# Traumatic Brain Injury Exacerbates Alzheimer’s Disease Pathology in the Retinas of TgF344-AD Rats

**DOI:** 10.1101/2021.09.23.461334

**Authors:** Conner Secora, Anne Vielle, Athena Ching-Jung Wang, Patricia Lenhart, Ernesto Salcedo, Noah R. Johnson, Md. Mahuiddin Ahmed, Heidi J. Chial, Timothy D. Boyd, Huntington Potter, M. Natalia Vergara

## Abstract

Alzheimer’s disease (AD) is a neurodegenerative condition that affects 6.2 million people age 65 and older in the U.S. alone, and is the leading cause of dementia. Moreover, AD can lead to visual impairment, and AD histopathology also manifests in the retina. However, the factors that modulate AD pathophysiology and lead to varied susceptibility and presentation in the population are not well understood. In this context, traumatic brain injury (TBI), which can arise from sport concussions, military combat, and other causes, is associated with a 2.3-fold higher risk of developing AD and AD-related dementias (ADRD). Thus, we set out to evaluate the effects of TBI, AD, and their combination, on retinal histopathology.

Several animal models have been developed to investigate the mechanisms underlying AD, but many have been limited by imperfect recapitulation of human pathology, and no model of TBI-associated AD (AD-TBI) has been characterized. To address this gap, we generated an innovative model of AD-TBI by taking advantage of a transgenic rat model (Tg-F344-AD) shown to recapitulate the main features of human AD pathology, and combining it with a two-time unilateral controlled cortical impact paradigm to mimic repetitive mild TBI (rmTBI). Histopathological analyses at four months post-impact confirm the presence of AD markers in transgenic retinas, and an increased severity of AD pathology due to TBI. Together, these results contribute to our understanding of the effects of TBI on AD retinopathy, with implications for patient care and therapeutic development.

## Introduction

More than 55 million people worldwide are living with dementia worldwide, and Alzheimer’s disease (AD) is the most common form of dementia accounting for 60-70% of all cases [1, 2]. Despite the high prevalence and devastating effects of AD, the factors that modulate the development of AD pathology and lead to different levels of severity and phenotypic presentation are incompletely understood. Among these, one factor that has been associated with AD is traumatic brain injury (TBI), which results from blunt or penetrative trauma to the head and affects an estimated 3.2-5.3 million Americans, with recent signs of increasing incidence rates [3, 4]. TBI is known to cause histopathological alterations that can ultimately lead to neurodegeneration and, in some cases, cognitive impairment [5, 6]. Epidemiological studies of TBI and AD have reported that individuals with a history of TBI have a 2.3-fold higher risk of developing AD and AD-related dementias (ADRD) [7]. Despite many investigations into AD and TBI individually, only a few studies have explored the correlations between the two pathologies, and particularly the long-term implications of their interaction in chronic conditions. Moreover, both TBI and AD have been associated with visual abnormalities that may be in part attributed to primary retinal histopathology [8–10].

The retina is the photosensitive region of the eye and is composed of neural tissue that originates from the neuroectoderm as an outcropping of the developing central nervous system (CNS) [11]. Cumulative evidence from the past decade suggests that the histopathological features of AD manifest not only in the brain, but also in the retina even prior to the onset of clinical symptoms [12]. The hallmark features of AD are amyloidosis and tauopathy, evidenced as amyloid-β (Aβ) plaques as well as neurofibrillary tangles (NFTs) that result from the abnormal aggregation of hyperphosphorylated tau proteins, typically found in the axons of neurons throughout the CNS [13, 14].

In this study, we investigated the correlation between TBI and AD and their long-term effects on retinal histopathology. In light of the limitations inherent to studies in human populations [15], there is a critical need for animal models that can faithfully recapitulate the pathogenesis of TBI and AD. Several mouse models have been used to investigate the pathophysiology of AD. However, mice have a natural resistance to tauopathy and do not recapitulate the full spectrum of AD histopathology seen in humans [9, 10]. We therefore sought to address this problem by capitalizing on a novel transgenic rat model of AD (i.e., TgF344-AD) that reliably mimics the main features of human AD pathology, including amyloidosis, tauopathy, and cognitive deficits, and combining it with a two-time, unilateral, closed-skull controlled cortical impact paradigm to establish an innovative model of repetitive mild TBI (rmTBI)-AD [10, 14]. Using the TgF344-AD model, we evaluated the impact of the interaction between rmTBI and AD on retinal thickness, retinal ganglion cell (RGC) numbers, cell death markers, neuroinflammation, and amyloid-β plaques in the retina of aged animals at four months post-injury, compared to control animals. Our findings show that the TgF344-AD rats show increased neuroinflammation and Aβ deposition in the retina, and that rmTBI enhances the severity of these phenotypes by increasing the formation of dense core amyloid plaques.

This study furthers our understanding of the role of TBI as a modulator of AD retinopathy, with potential implications for patient care and therapeutic interventions.

## Materials and Methods

### Rat Model of AD-TBI

Animal handling and experimentation was approved by the Animal Care and Use Committee of the University of Colorado. This study was performed using TgF344-AD rats (from the PrP-hAPP[K670N/M671L]/PrP-hPSEN1[dE9] strain developed on a Fischer 344 [F344] rat background). These transgenic rats harbor mutant human genes for APP (with the Swedish mutations at amino acids 595/596) and presenilin 1 (PSEN1, deltaE9) [10, 16]. Wild-type (WT) F344 rats were used as controls.

At 12 months of age, half of each of the two cohorts (wild-type and TgF344-AD) were subjected to two events (separated by one week) of unilateral closed-head controlled cortical impact (2xCCI) using a stereotaxic apparatus with a flat 5 mm diameter metal tip attached to a Leica Impact One device, delivering an impact at 6 m/s with parameters previously reported for intact scalp CCI [3]. The 2xCCI were administered to isoflurane-anesthetized rats, at an impact site between the bregma and lambda, 3 mm to the right from the rat’s midline. The other half of the cohort underwent a “sham injury” that consisted of an identical protocol for the 2xCCI minus the actual impact. This experimental design resulted in four groups for analysis: wild-type-sham (WT-S), wild-type with 2xCCI (WT-TBI), transgenic-sham (AD-S), and transgenic with 2xCCI (AD-TBI).

Animals were euthanized at 120 days post-injury by overdose of sodium pentobarbital (100 or greater mg/kg IP), perfused with 0.9% saline, and organs were harvested for post-mortem pathological analyses. For this study, the left eye of each rat was enucleated and prepped for histological examination after immersion in Davidson’s fixative for 24 h.

### Histological Methods

Eyes were dehydrated through immersion in ethanol of increasing concentrations followed by xylene, embedded in paraffin, and sectioned at 5 μm thickness using a Leica microtome. Sections at regularly occurring intervals were stained using hematoxylin and eosin to analyze the general morphology of each eye and to determine key anatomical landmarks. For each eye, four regions of interest (ROIs) of the posterior retina were defined using the optic nerve as a landmark, with each region consisting of a 250 μm stretch of retina approximately 700 μm in distance from the optic disc, one in each of the four quadrants of the retina. These ROIs were then tested for markers of inflammation, neurodegeneration, and AD pathology using immunohistochemistry as follows: sections were rehydrated, and antigen-retrieval was performed using 1x citrate buffer for 7 min in an autoclave at approximately 125°C. Sections were then incubated with blocking solution consisting of 10% normal donkey serum (NDS) diluted in 1xPBS (0.25% Triton), followed by incubation with primary antibodies (see Table1) overnight at 4°C. Incubation with secondary antibodies (see Table 1) was performed for 1 h at room temperature. All antibodies were diluted with 2% NDS in 1xPBS (0.25% Triton). A fluorescent amyloid-binding dye, NIAD-4 (diluted in 1xPBS), was used as an additional marker for Aβ deposits. A Nikon Ti Eclipse microscope was used for fluorescent confocal imaging, and a Nikon Ti2 Eclipse WideField microscope was used for brightfield imaging.

**Table 1.**

|  | **Reagent** | **Dilution** | **Manufacturer** | **Catalog #** |
| --- | --- | --- | --- | --- |
| Primary Antibodies | Mouse anti-amyloidβ 4G8 - monoclonal | 1:200 | BioLegend | 39220 |
|  | Mouse anti-GFAP - monoclonal | 1:500 | Proteintech | 60190-1-Ig |
|  | Mouse anti-HuC/D - monoclonal | 1:100 | Invitrogen | 16A11 |
|  | Rabbit anti-CC3 - monoclonal | 1:250 | Cell Signaling Technology | 5A1E |
| Secondary Antibodies | AlexaFluor 488-conjugated donkey anti-mouse IgG | 1:1,500 | Invitrogen | A32766 |
|  | AlexaFluor 546-conjugated donkey anti-mouse IgG | 1:2,000 | Invitrogen | A10036 |
|  | AlexaFluor 488-conjugated donkey anti-rabbit IgG | 1:1,000 | Invitrogen | A-21206 |
| Fluorescent Dye | NIAD-4 | 1:500 | Cayman | 18520 |

### Image Analysis and Statistics

The retinal layers were measured using the Nikon Elements imaging software. Analyses of fluorescence micrographs were automated using MATLAB (version R2021a, product of MathWorks). The data for each ROI were added as a representation of each eye, and then compiled into their respective groups for statistical analyses using Prism (version 8.4.3, product of GraphPad Software, LLC). All data sets were normally distributed and compared using oneway analysis of variances (ANOVA) with Tukey post-hoc multiple comparisons test (α = 0.05). In all graphical representations, error bars represent the standard error of the mean (SEM).

## Results

### Morphometric Analyses of the Retina and Retinal Ganglion Cell Count

The morphology of the retina was assessed on hematoxylin and eosin-stained retinal sections from animals in each of the four study groups (WT-S, WT-TBI, AD-S, and AD-TBI; n=5 animals per condition) to determine whether there were any observable changes due to tissue atrophy. Previous studies have reported significant atrophy across multiple layers of the retina in postmortem samples from humans with AD, along with other animal models including the TgF344-AD rat model used in this study [2, 9, 17]. Here, measurements of total and individual retinal layer thicknesses were obtained for each eye (individual layer thicknesses were reported as fractions of the total retina). Measurements for both the nerve fiber layer (NFL) and the retinal ganglion cell (RGC) layer were combined due to the difficulty in delineating the boundary between the two layers. Notably, no significant differences in the thicknesses of the total retina or in the thicknesses of any of the individual layers of the retina were detected in this study (Fig. 1B-E, K) or in CC3 levels (Fig. 1L) among the four groups. Together, these results suggest a low level, if any, of neurodegeneration within the retina at the selected time point. Although surprising, some reports have noted a similar difficulty in detecting differences in RGC numbers in this rat model [9]. It is possible that this finding may also simply reflect the current study’s low power for detecting significant differences in RGC numbers, if present. Notably, the majority of CC3 signal was found in blood vessels within the NFL (not shown). This may be an indication of vascular compromise due to CCI injury, or of amyloidosis angiopathy, which is often associated with cerebral AD pathology and may contribute to early stages of neurodegeneration [20, 21]. Further studies are needed to test these hypotheses.

**Figure 1.**
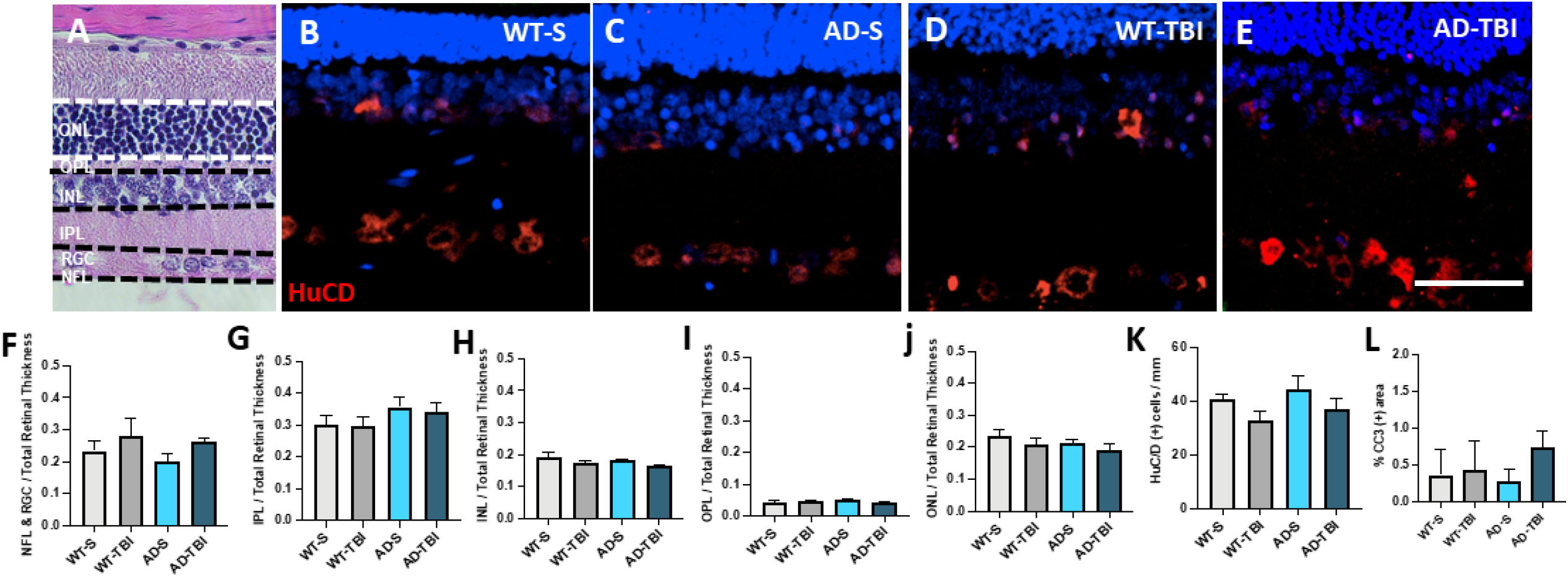
No significant differences in retinal thickness, RGC count, or CC3 expression were detected in TgF344-AD rats with or without rmTBI treatment compared to controls. **(A)** Individual retinal layers were measured on H&E stained sections and reported as a fraction of the total retinal thickness. **(B-E)** Representative micrographs of HuC/D (red) immunostaining in the retina of the four different groups: WT-S, AD-S, WT-TBI, and AD-TBI. Scale bar = 50 μm. **(F-J)** No significant differences in individual retinal layer thickness were observed among groups (n = 4 animals per group, analysis performed with ANOVA, α = 0.05). **(K-L)** No significant differences were identified among groups in the numbers of HuC/D positive cells per mm of retina (**K**; n=3, p=0.34) or in the area of CC3 staining (**L**; n=4, p=0.23). Error bars represent S.E.M. ONL: outer nuclear layer; OPL: outer plexiform layer; INL: inner nuclear layer; IPL: inner plexiform layer; GCL: ganglion cell layer; NFL: nerve fiber layer.

### AD Retinas Exhibit Increased Neuroinflammation

Inflammation is one of the hallmark features of brain AD pathology [12, 13, 22], and it is also present to varying degrees during both the acute and chronic phases of TBI [3, 5]. To evaluate the independent and potentially synergistic contributions of AD and TBI on retinal neuroinflammation, we examined glial fibrillary acidic protein (GFAP) expression throughout the retina. GFAP is an intermediate filament expressed in astrocytes, and its expression levels increase significantly in Müller glia cells during reactive gliosis as part of an inflammatory process within the retina [6, 22]. No differences were observed in GFAP expression in the NFL across all groups (Figure 2E). However, the AD groups exhibited significantly higher levels of GFAP expression in Müller glia compared to WT-S controls (Figure 2F), suggesting that AD induces retinal neuroinflammation with activation of Müller glial cells. Although not statistically significant in our study, TBI alone was associated with a trend towards increased GFAP reactivity in WT animals. However, once the neuroinflammatory process is established in the AD retina, TBI may not be able to further increase the Müller glia response above baseline.

**Figure 2.**
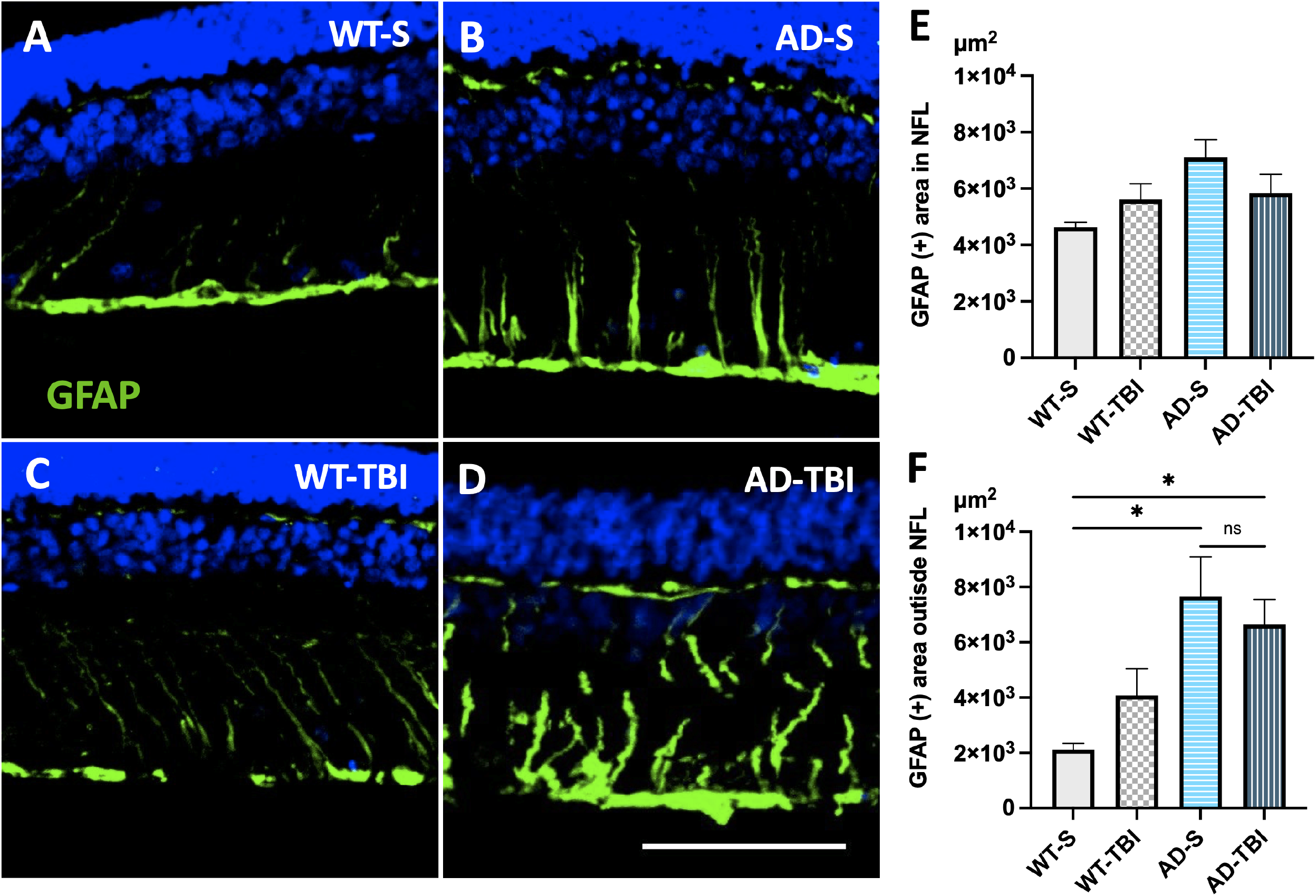
TgF344-AD rat retinas display increased Müller glia activation. **(A-D)** Representative micrographs show GFAP expression patterns in the retinas of WT-S, AD-S, WT-TBI, and AD-TBI rats, respectively. Scale bar = 50 μm. **(E)** No significant changes in GFAP expression in the NFL, which encompasses the astrocytic population, were detected among the four groups. **(F)** Tgf344-AD transgenic groups display increased GFAP immunoreactivity in the retina compared to the WT groups (n = 5, analyzed using ANOVA p=0.0088; * p<0.05). Error bars represent S.E.M.

### TBI Increases the Severity of Aβ Pathology in AD Retinas

There is mounting evidence for the amyloidosis-driven model of AD pathogenesis, which is supported by the full spectrum of AD pathology reported in the TgF344-AD rats that only express mutant human genes that induce the accumulation of Aβ peptides [10, 13, 16]. Considering this growing body of evidence, we sought to quantify Aβ peptide aggregates using the 4G8 antibody (with affinity to amino acids 17-24 of Aβ) along with the fluorescent dye NIAD-4 to assess Aβ plaque formation. NIAD-4 has a high affinity for Aβ fibrils and displays increased fluorescence intensity when bound to β-pleated sheet structures, which are found in dense core Aβ plaques [23, 24]. As expected, significant numbers of Aβ peptide aggregates were found in both AD groups compared to WT (Figure 3). Even though AD rats subjected to 2xCCI showed a trend towards increased numbers of Aβ aggregates compared to the AD-S rats, the difference was not statistically significant in our study. Notably, Aβ aggregates stained by both the 4G8 antibody and NIAD-4, suggestive of dense core plaque formation, were only present to a significant degree in AD-TBI rats compared to all other groups (Figure 4). This finding supports the hypothesis that TBI increases the severity of AD pathology in the retina.

**Figure 3.**
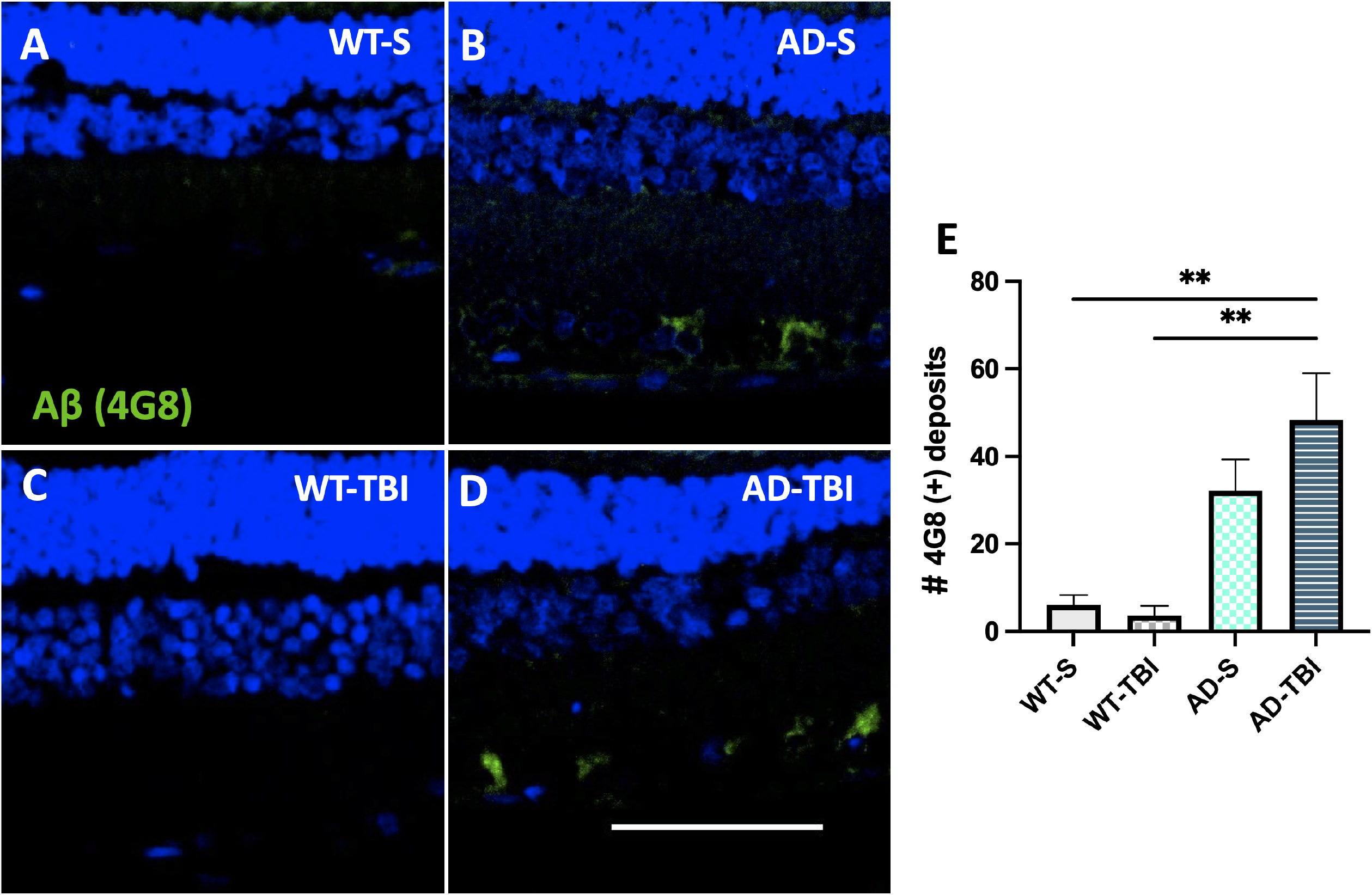
Increase in the numbers of Aβ aggregate deposits in the retinas of TgF344-AD rats. **(A-D)** Representative micrographs show patterns of 4G8 antibody (green) immunostaining, indicative of Aβ peptide deposition, in the RGC layer of the four different groups: WT-S, AD-S, WT-TBI, and AD-TBI. Scale bar = 50 μm. **(E)** Quantification of Aβ aggregates shows a marked increase in Aβ deposition in both transgenic TgF344-AD groups (n=5; ANOVA with Tukey-modified post-hoc comparisons; ** p<0.01). Error bars represent S.E.M.

**Figure 4.**
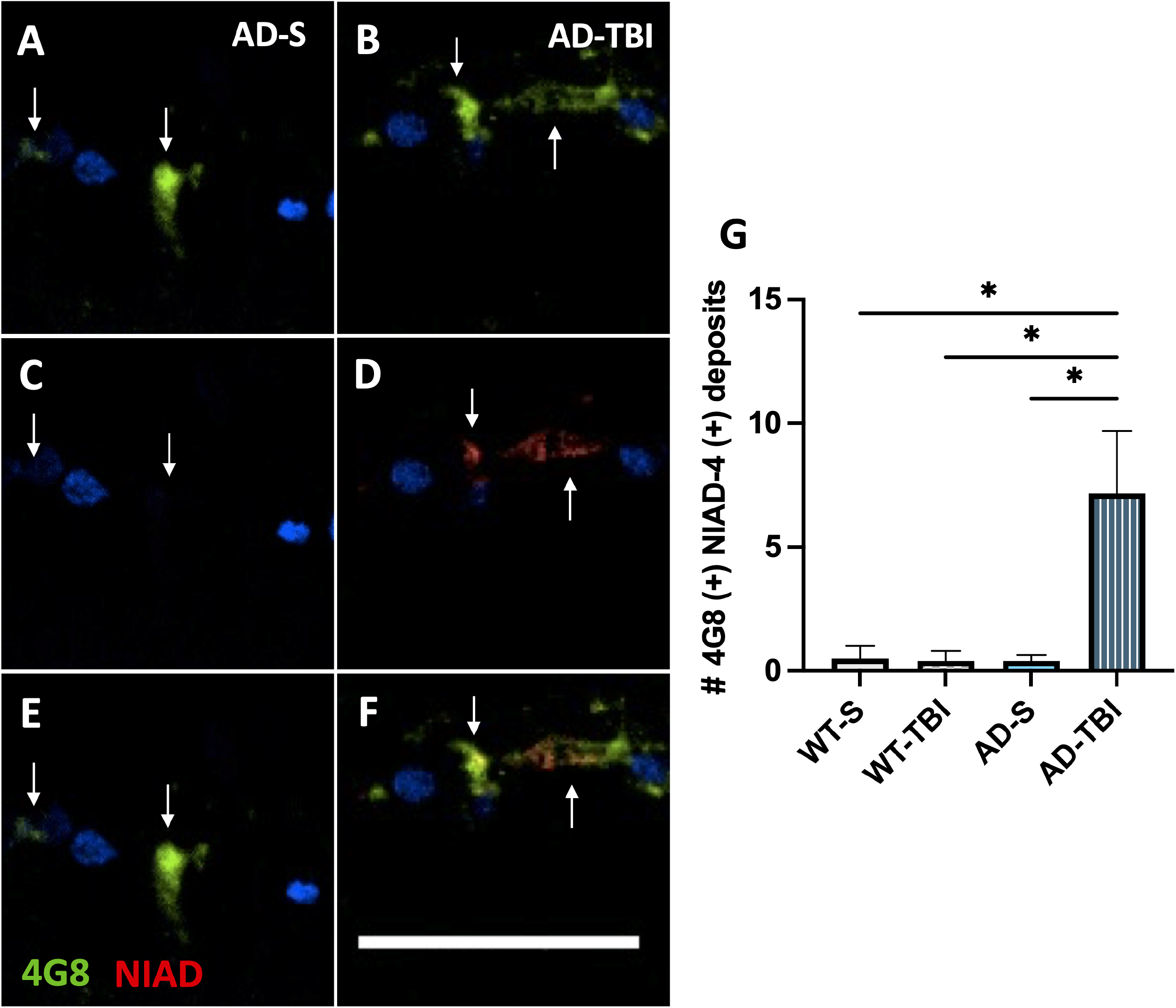
TBI increases the formation of dense core amyloid plaques in the retinas of TgF344-AD rats. **(A-F)** Representative micrographs of 4G8 (green) and NIAD-4 double-staining in the RGC layer of AD-Sham (A,C,E) and AD-TBI (B,D,F) animals shows colocalization of these markers only in AD-TBI group, indicating amyloid plaque maturation. Scale bar = 50 μm. **(G)** Quantification of 4G8(+) NIAD-4(+) deposits shows a significant increase in amyloid plaques in AD-TBI rat retinas compared to all other groups (n=5, ANOVA with Tukey-modified post-hoc comparisons, * p<0.05). Error bars represent S.E.M.

## Discussion

Individual explorations of TBI and AD have revealed processes of neuroinflammation and vascular injury that ultimately lead to neuronal loss in both conditions, with associated visual dysfunction. However, this is the first time that TBI has been characterized in the TgF344-AD rat model, linking the two conditions in an animal model that can recapitulate the full spectrum of human AD pathology. This study did not detect any significant evidence of neurodegeneration within the retina evidenced by reduction in retinal thickness or RGC numbers, which contrasts with reports of human pathology as well as with studies of animal models, including the TgF344-AD rat [2, 9, 12, 17, 25]. However, the rodent models cited in other reports were typically older than 16 months, which could indicate that the rats used in our study were still in the earlier stages of AD pathophysiology in the retina. The localization of CC3 in blood vessel walls could be suggestive of amyloidosis angiopathy or of persistent vascular compromise due to TBI. Vascular abnormalities due to pericyte and endothelial cell death wrought by cerebral amyloidosis angiopathy have been noted to lead to subsequent tissue injury and are thought to be a critical component in the disruption of the blood-brain barrier (BBB) and the reduction of Aβ clearing mechanisms, leading to an overall worsening of amyloidosis [20, 26, 27]. Thus, we raise the possibility that this may be the case in the retina as well. Additional studies focused on the retinal vasculature are needed to elucidate the role of amyloidosis angiopathy in the retina and its relationship to TBI.

Despite the absence of significant differences in neurodegeneration reported in this study, the AD retinas did show signs of increased neuroinflammation. The extension of GFAP into the outer layers of the retina observed in the AD groups indicate a heightened glial response mediated by Müller cells, which is typically seen in advanced stages of retinal inflammation [6, 22]. The role of inflammation in response to injury is an area of ongoing investigation and has been shown to be a complex, dynamic response that involves locally recruited glial mechanisms in addition to immune and vascular mechanisms that manifest both locally and systemically [3, 21, 22]. This study only tested an intermediate component of the larger processes at work during chronic inflammation. Patterns of reciprocal induction that regulate inflammation have been described between microglia, astrocytes, and circulating leukocytes [28]. A more comprehensive examination of resident immune cells and local and systemic inflammatory markers would elucidate potential differences in both the magnitude and phenotypes of chronic inflammation observed in this animal model.

Moreover, this study shows for the first time a significant increase in dense Aβ plaque formation in AD retinas exposed to TBI. This supports the hypothesis that injury caused by TBI can exacerbate AD retinal pathology, and merits further investigation into the underlying mechanisms of AD pathology as they relate to TBI.

In summary, we have established an innovative rat model of AD-TBI as a useful tool to evaluate the relationship between TBI and AD in the retina. This model will be critical in expanding the body of knowledge concerning the chronic, long-term effects of TBI, which remain poorly understood [3, 4, 29]. Further characterization of this animal model will facilitate the elucidation of the many interconnected disease mechanisms involved in AD and TBI, which is likely to improve risk assessment and therapeutic development for individuals who are predisposed to developing AD. The characterization of retinal pathology may also help further the development of the retina as a biomarker for the early detection of AD pathology before the onset of clinical symptoms [30]. Ultimately, this may contribute to much-needed advances in clinical assessment and treatments for individuals suffering from AD and/or TBI.

## Acknowledgements

This work was supported in part by a Challenge Grant to the Department of Ophthalmology at the University of Colorado from Research to Prevent Blindness, by the Linda Crnic Institute for Down Syndrome, and by DoD grant W81XWH-17-1-0583 to H.P.

